# An Open-Source Neurodynamic Model of the Bladder

**DOI:** 10.1101/2024.11.21.624716

**Authors:** Elliot Lister, Aidan McConnell-Trevillion, Milad Jabbari, Abbas Erfanian, Kianoush Nazarpour

## Abstract

**Background:** Lower urinary tract (LUT) symptoms affect a significant proportion of the population. In-silico medicine can help understand these conditions and develop treatments. However, current LUT computational models are closed source, too deterministic, and do not allow for simple use of modelling neural intervention.

**Methods:** An open-source Python-based model was developed to simulate bladder, sphincter, and kidney dynamics using normalised neural signals to predict pressure and volume. The model was verified against animal bladder data, assessed for noise sensitivity, and evaluated against known physiological factors.

**Results:** The animal data comparison yielded a significantly more similar pattern than existing models, with a correlation coefficient of *r* = 0.93 (*p* < 0.001). All physiological factors were within bounds, and the model remained stable with noise under the described boundaries.

**Conclusion:** The proposed model advances the field of computational medicine by providing an open-source model for researchers and developers. It improves upon existing models by being accessible, including a built-in neural model that better replicates smooth bladder filling results, and incorporating a novel kidney function that alters bladder function by time of day in line with circadian rhythm. Future applications include personalised medicine, treating LUT symptoms with in-silico models and adaptive neural interventions.

## 1 Introduction

Lower urinary tract (LUT) symptoms are a diverse and highly prevalent set of conditions known to affect over 60% of adults by the age of 40 [1]. Overactive bladder, a type of LUT symptom [2], is particularly widespread - with a reported prevalence of 10.7% of the global population [3] and 12% in the United Kingdom alone [4]. Overactive bladder is typically characterised by urinary urgency, frequency, and urge incontinence [4] and can impart significant impacts on quality of life, causing social embarrassment, sleep disruption, and anxiety [5]. Despite the availability of effective treatments, such as bladder retraining and pharmaceutical intervention [6], limited knowledge about the underlying aetiology remains a significant barrier to the development of more efficient solutions with fewer side effects. This research gap may be addressed by the development of computational models of the LUT. Computational approaches to LUT research offer a number of benefits. Short-term benefits of these models include increased reproducibility, which allows researchers to share and validate their findings. Additionally, these models can contribute to a reduction in animal testing, promoting more ethical and efficient research practices. In the long term, clinicians and researchers may utilise simulated organs to integrate individual patient characteristics and unique disease states. This personalised approach enables the prediction of treatment outcomes under different scenarios, guiding the selection of the most effective treatment strategy. By minimising time spent trialling ineffective treatment plans, valuable time and resources can be conserved. This benefit is particularly enticing given the stark economic cost of overactive bladder, with £840 million spent annually in the UK on the treatment and management of the condition [7]. Moreover, models play a considerable role in the design and optimisation of novel medical devices. Despite the clear benefits of a computational approach, present attempts at capturing the system fall short on several fronts.

A robust model of the LUT requires several key attributes. Firstly, the capacity to integrate external neural inputs is essential for capturing complex interactions with the nervous system. Secondly, a smooth neural model representation is essential to avoid unrealistic action potentials. Thirdly, incorporating variable kidney function is fundamental for preventing oversimplified and deterministic outcomes. Lastly, code accessibility is necessary to facilitate model sharing, validation, and broader application within the scientific community.

Current LUT models, see Table 1, use unrealistic representations of biological processes, such as treating urine inflow from the kidneys as a constant rate. These assumptions may lead the model to deviate from physiological observations, thereby impeding its ability to accurately represent LUT behaviour. Moreover, some models suffer from problems related to reproducibility. The original papers may not provide enough detail for accurate replication, thereby creating obstacles for other researchers to effectively utilise and advance these models, consequently impeding scientific progress.

**Table 1:** Comparison of existing models and their parameters.

| Model | Inputs | Neural model | Kidney function | Outputs | Code availability |
| --- | --- | --- | --- | --- | --- |
| **Our model** | Neural units for detrusor activation, detrusor inhibition, and sphincter activation. | Piecewise Gaussian | Stochastic | Pressure and volume by unit time | Open access |
| Valentini, Besson, and Nelson (VBN) [8] | Gender, initial bladder volume, detrusor and sphincter activity, and urethral properties. | Step function with decay components. | N/A | Flow rates and bladder pressure over time | Upon request |
| Hosein and Griffiths [9] | N/A | Threshold-based, firing-rate model with afferent feedback | Constant | Pressure and volume by unit time | Upon request, but in legacy Pascal. |
| Fletcher [10] | Neural units for bladder musculature and urethral relaxation. | Exponential | N/A | Pressure and volume by unit time | No |
| Paya *et al.* [11] | N/A | Neural Regulator System | Constant | Pressure and volume by unit time | No |
| Bastiaanssen *et al.* [12] | Neural units for detrusor activation, detrusor inhibition, and sphincter activation. | Manually tuned, firing-rate neural network [13] | Constant | Pressure and volume by unit time | No |

This paper aims to overcome these limitations by introducing a novel and rigorously validated computational model of the LUT. Our model incorporates several significant modifications. Firstly, it features a dynamic urine inflow that reflects the body’s natural fluctuations in urine production throughout the day, mimicking natural physiological patterns for a more accurate representation of bladder filling. Secondly, the model incorporates elements of randomness to account for the inherent variability present in biological systems. The inclusion of stochasticity in the model enhances its realism, ensuring that it accurately represents the full range of bladder behaviour while operating within safe limits. Thirdly, we use Gaussian functions to model neural inputs from the brain, resulting in a more gradual rise in bladder pressure compared to existing models that utilise step-like functions. Finally, recognising the importance of collaboration and knowledge sharing in scientific progress, we have made the code associated with this model open access for other researchers to utilise and build upon.

Figure 1 illustrates the biological system (left) and its computational analogue (right). Three normalised neural inputs modulate the detrusor and sphincter muscles, thereby controlling outflow rate. Urine inflow, modelled independently by a kidney function, determines bladder filling. A feedback loop incorporates bladder pressure to dynamically update the neural model.

**Figure 1:**
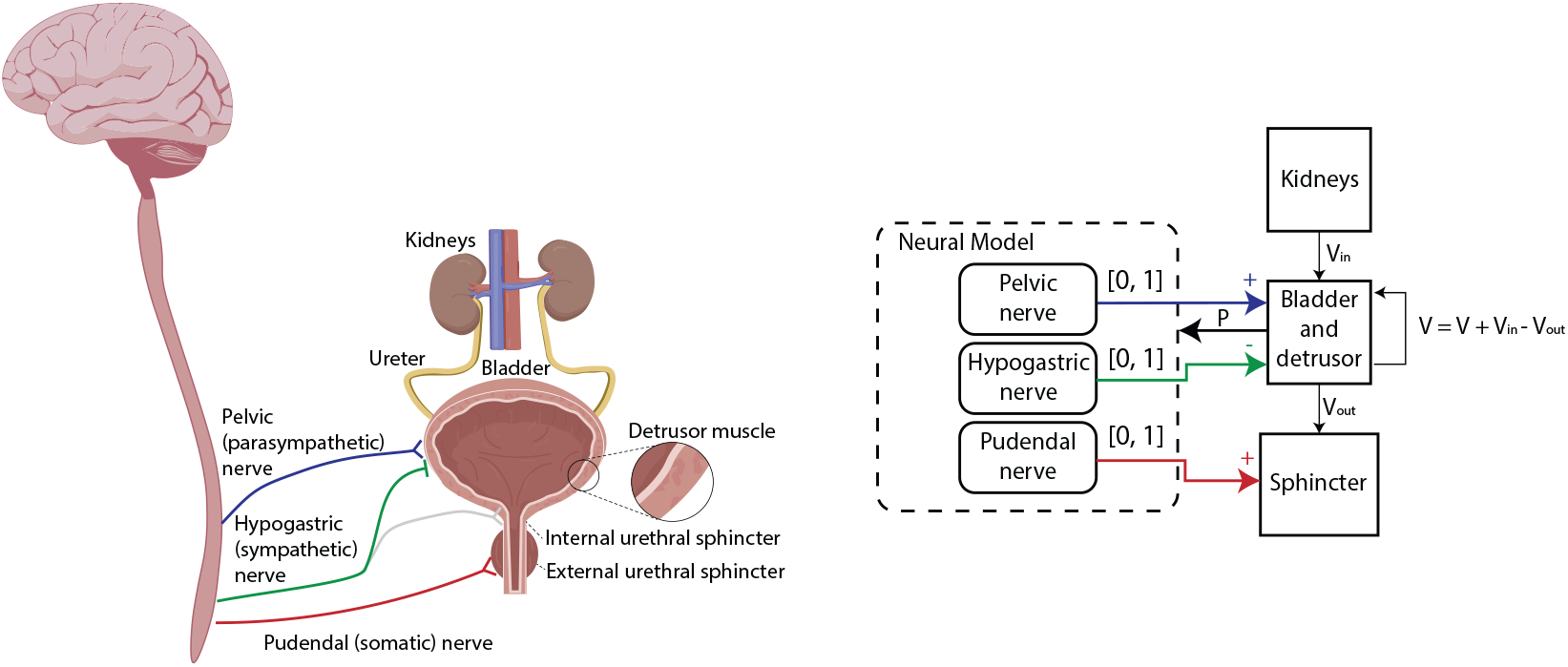
Left: Anatomy of the lower urinary tract and kidney, highlighting the detrusor muscle, urethral sphincters, and their innervation. Right: Computational analogue of the LUT system.

This comprehensive and accessible model paves the way for the development and testing of in silico interventions. Through offering a more realistic portrayal of bladder function, our model can have a significant impact in multiple domains.

## 2. Method

### 2.1 Core Bladder Function

Our model is inspired by the framework developed by Bastiaanssen *et al*. (1996) [12]. It uses three normalised inputs to represent the neural control of the bladder and sphincter: 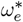 (excitatory input of the detrusor muscle), 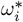 (inhibitory input of the detrusor muscle), and 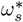 (excitatory input of the sphincter), corresponding to the pelvic, hypogastric, and pudendal nerves, respectively. The model focuses on three state variables, namely,

- *V*_*B*_: The volume within the bladder.
- 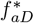: The normalised activation of the detrusor muscle, and
- 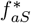 : The normalised activation of the sphincter muscle.

Each state variable is modelled and updated by a differential equation, where the neural activations make use of time constants *t*_*D*_ and *t*_*S*_. We refer to the functions that model each equation as *f*_1_, *f*_2_, and *f*_3_, respectively:

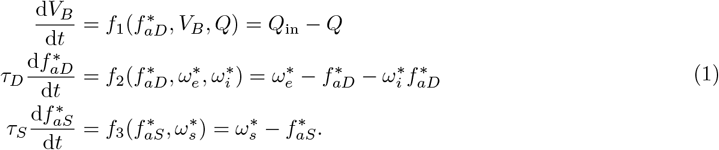

Here, *Q*_in_ is the urine inflow determined by the kidney model. *Q* is the urine outflow, which is a function of the urethral opening radius *r*_*U*_, determined using a mapping function *f*_map_ based on *V*_*B*_, 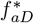, and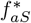. The root of the function 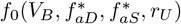 is found using the bisection method, implemented with the SciPy package [14]. If the bisection method fails, *r*_*U*_ defaults to zero, assuming the urethra remains closed.

Muscle tension in the sphincter and detrusor includes both active and passive components. Passive tension is modeled by an exponential relationship dependent on the urethral radius *r*_*U*_ :

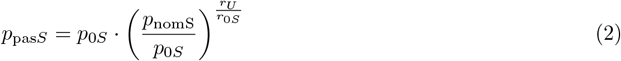

Active pressure within the urethra depends on the muscle tension in the sphincter wall (*σ*_*S*_) and the curvature of the sphincter. Given the assumption that the sphincter muscle can be modelled as a cylinder, it exhibits unidirectional bending. Hence, the active pressure can be represented by the equation:

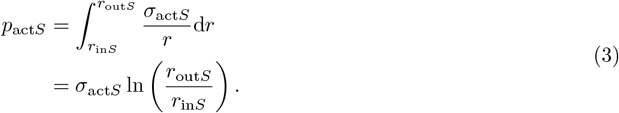

The active tensile stress *σ*_nom_act*S*_ is calculated based on the normalized activation 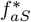, isometric stress *σ*_isoS_, and the force-velocity relationship 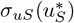:

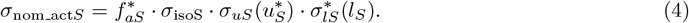

The contraction velocity *u*_*S*_ is determined by the rate of change of the sphincter radius *r*_*S*_:

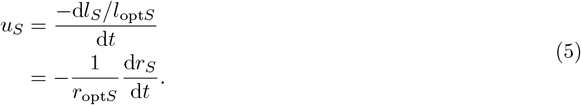

The rate of change of *r*_*S*_ is given by:

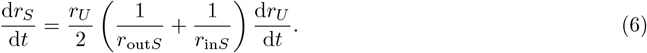

Thus, the contraction velocity of the sphincter muscle is:

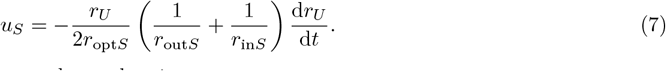

Hence, the pressure in the sphincter muscle can be given as:

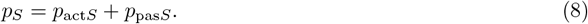

Bastiaanssen *et al*. [12] derived a relation between the urethral radius (*r*_*U*_) to the square rate of urine flow (*Q*^2^) as

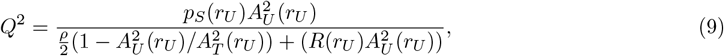

where *A*_*U*_ is the cross-sectional area occupied by urine. The detrusor muscle pressure calculation follows a similar approach, with the contraction speed *u*_*D*_ given by:

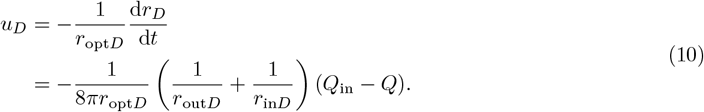

Assuming *Q*_in_ = 0, the detrusor muscle tensile stress includes elastic and visco-elastic elements of the bladder wall. The objective function (*f*_0_) calculates the pressure difference between the bladder neck and the urethra entrance, considering factors like bladder volume (*V*_*B*_), muscle contraction strength 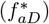, sphincter resistance 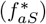, and urethral radius (*r*_*U*_). The bladder volume (*V*_*B*_) is updated based on the calculated net flow (*f*_1_) through the use of the following equation:

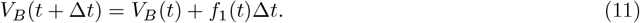

For detailed derivations and a thorough explanation of the underlying formulas, readers are referred to the original paper [12].

### 2.2 Neural Model

The neural control model used with the bladder model [13] could not be replicated due to insufficient data and descriptions of fitting techniques. Therefore, we developed a new model using piecewise Gaussian functions to simulate the neural inputs. Each input is modeled using a left-skewed Gaussian function, where the slope is smooth until the point of voiding, at which the inputs are fixed as constants. The bladder excitatory 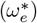 and inhibitory 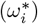 inputs are functions of the bladder volume (*V*_*B*_). These inputs provide feedback based on the current volume of the bladder. The functions are defined as follows:

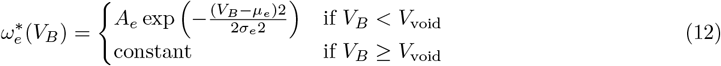

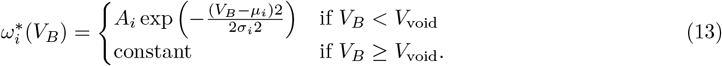

Here, *A*_*e*_ and *A*_*i*_ are the amplitudes, *µ*_*e*_ and *µ*_*i*_ are the means, and *s*_*e*_ and *s*_*i*_ are the standard deviations of the Gaussian functions. *V*_void_ is the bladder volume at which voiding occurs, and beyond this point, the inputs are fixed as constants. The sphincter muscle excitatory input 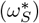 is a function of the urethral radius (*r*_*U*_). This input provides feedback based on the current radius of the urethra. The function is defined as follows:

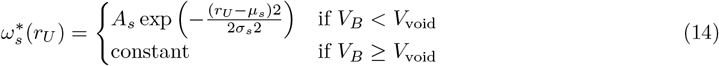

Similarly, *A*_*s*_ is the amplitude, *µ*_*s*_ is the mean, and *σ*_*s*_ is the standard deviation of the Gaussian function. This piecewise Gaussian approach allows for smooth transitions in neural inputs until the point of voiding, ensuring realistic simulation of bladder and sphincter muscle control.

### 2.3 Kidney Function

Modelling the state of kidney outflow for an individual is challenging because of factors such as liquid intake, food consumption, medications, other medical conditions, age, and perspiratory behaviour [15]. Given the number of parameters involved, it would not be feasible to implement or validate a kidney model relying on these factors. However, there is one factor that universally affects bladders: time. Although the production of urine in the kidneys does not change intensely throughout the day, it has been shown that numerous functions of the kidney, including urine production, vary with circadian rhythm [16, 17]. Based on this information, the inflow (*Q*_in_) has been represented mathematically as a sinusoidal function.

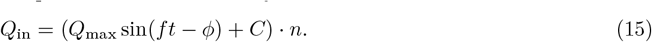

Where *Q*_max_ is the max inflow, *f* is the urinary frequency, *t* the simulation time, *ϕ* the phase shift, *C* a constant term, and *n* is the uniform noise. After each minute (or chosen time step), a random scalar (*n*) uniformly distributed between -1 and 1 is multiplied by the volume obtained from the sine function. This method preserves the overall circadian pattern while introducing controlled variability. As insufficient public data exists on per minute urine production, the function was fit to align with an average inflow that experiences fluctuations over a 24-hour cycle within ranges of typical urine production. With that in mind, the numbers provided can be adjusted and personalised, thereby enhancing the model’s compatibility with users. Using this function creates peaks in urine production during the afternoon and weaker inflow during the early morning, aligning with what is reported in the literature [16]. Despite its increased complexity, this model of urine production can still be adjusted with just a few parameters if there were a significant difference between a patient’s flow rates and the current model fit.

### 2.4 Code Availability

The code for implementing the bladder model, neural control model, and kidney function can be accessed via the GitHub repository [18]. Access to the repository is open for researchers and developers who are interested in reviewing, replicating, and building on the work presented in this study. The repository provides detailed documentation and examples, which should support users in understanding and utilising the content.

### 2.5 Validation

Validating a biophysical LUT model is inherently challenging due to the invasive nature of acquiring high-quality clinical data on bladder pressure, volume, and voiding dynamics. Ethical constraints limit experiments on healthy individuals, and variations in anatomy and physiology among patients complicate direct comparisons between model predictions and clinical observations. Additionally, privacy laws often restrict access to human data, hindering reproducibility. To address these challenges, we have validated model components using data from animal studies and published physiological ranges of human data. This involved comparing model outputs to established limits, reproducing published results, and analysing model behaviour under known physiological perturbations. Utilising data previously obtained from animal experiments [19], a two-step validation was employed to assess the lower urinary tract model’s ability to represent real-world physiology. First, min-max scaling was applied to the animal data to bring it into a range comparable with the predictions of the human model. This allowed for a visual overlay of the scaled animal data onto the model’s outputs. This qualitative comparison provided valuable insights into the similarity between the model’s behaviour and the actual observations from the animal experiments. Secondly, a quantitative analysis using Pearson correlation was conducted. In this case, the ratio of pressure to volume for the non-scaled animal data was compared with the corresponding outputs from the human model.

## 3. Results

This section reports the core outputs and dynamics of the proposed bladder model. We explore the pressure-volume relationship and compare our simulations to experimental animal data. Additionally, we examine key physiological metrics such as voiding durations and filling times. Finally, we assess the robustness of the model to noise perturbations.

### 3.1 Model Behaviour

Figure 2 illustrates the dynamic behaviour of the model, showcasing the time-series evolution of bladder pressure, volume, and neural activity. The plot demonstrates the expected cyclic nature of bladder function, alternating between storage and voiding phases. During the storage phase, bladder volume gradually increases in a stochastic manner, reflecting the random inflow from the kidneys. As the bladder fills, neural activity gradually increases, modelled by a piecewise Gaussian function. To simulate the rapid transitions during the voiding phase, a simplified method was utilised. In this method, the neural units remain constant during voiding, as depicted in Figure 2, leading to a stepwise alteration in neural activity. This simplification allows for easier manipulation of voiding dynamics, enabling the exploration of various scenarios and the tuning of model behaviour (e.g. voiding speed). For example, increasing the sphincter neural input 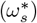 from 0.1 to 0.25 during voiding can significantly prolong voiding time, from 27.1s to 308.6s. Conversely, decreasing this input to 0.01 can shorten voiding time to 25.4s. Similarly, increasing the excitatory input to the bladder 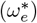 from 0.6 to 1.0 can, as expected, reduce voiding time to 20.2s due to the increased bladder pressure. These simulations highlight the model’s sensitivity to parameter changes and its ability to capture a range of physiological behaviours. While careful consideration should be given when modifying these values to avoid unintended consequences, this flexibility is a key advantage of the model, allowing for the exploration of various neurotherapeutic strategies and the development of innovative neuromodulation techniques. In the subsequent subsections, the neural model parameters will be reset to their original values, and further tests will be performed to validate the LUT model.

**Figure 2:**
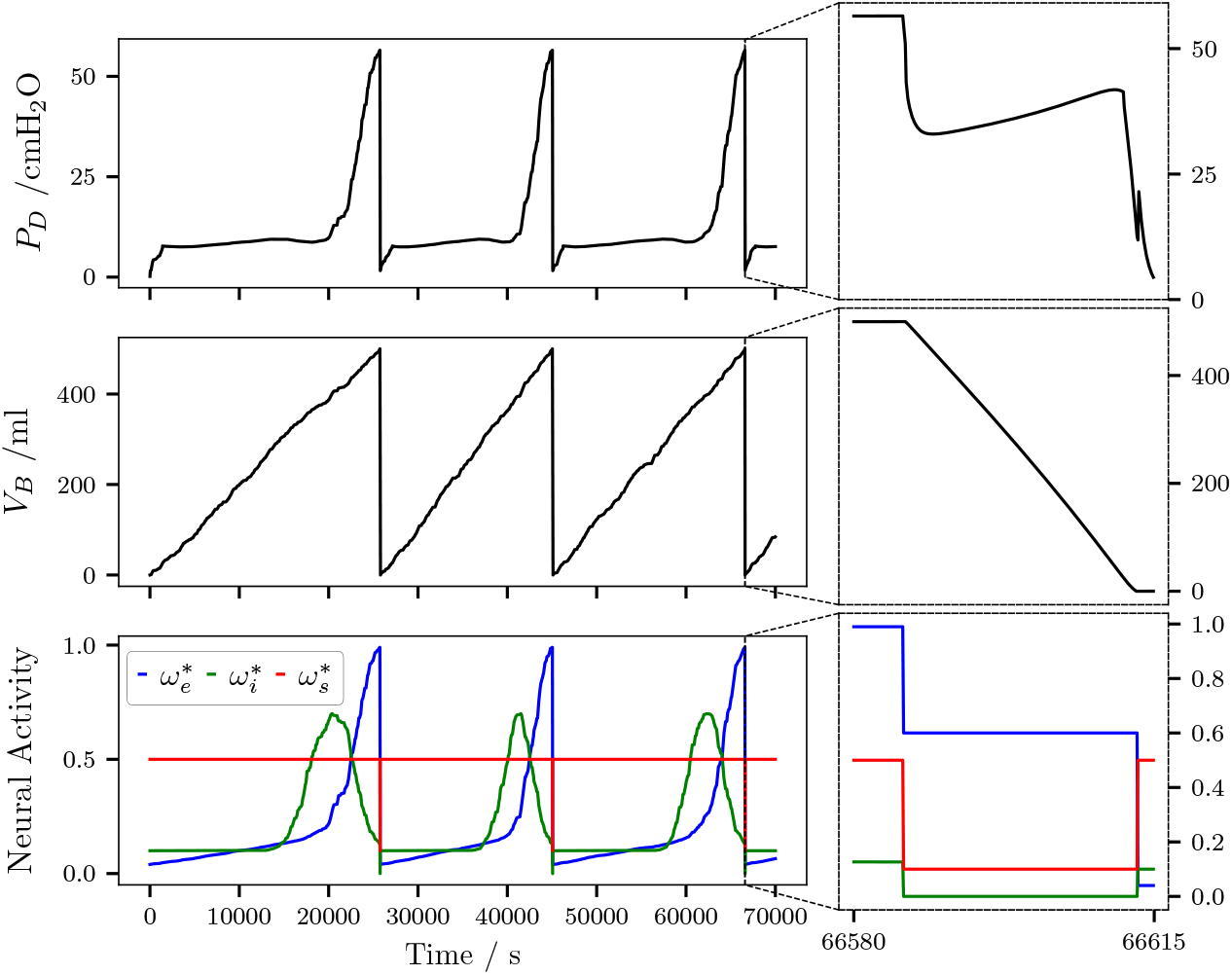
Time series plots illustrating the dynamic behaviour of the model, including bladder pressure, volume, and neural activity. A zoomed-in view of the voiding phase is also provided to highlight the rapid changes in these variables during micturition. *V*_*B*_: Bladder volume, *P*_*D*_: Detrusor pressure, 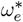: Excitatory bladder neural input, 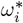: Inhibitory bladder neural input, 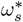: Sphincter neural input.

### 3.2 Pressure-Volume Relationship

In contrast to Bastiaanssen’s neural model [13], our model results in a slower increase in pressure. This modification in the neural weights’ interaction with the model leads to a less pronounced slope in the pressure-volume relationship. To facilitate comparison with literature, the detrusor pressure has been converted to units of cmH_2_O. The pressure-volume plot in Figure 3 exhibits a close resemblance to an ideal cystometrogram of a typical adult human female [20], characterised by an initial rise in pressure during phase I, subsequent bladder compliance in phase II, and concluding with a gradual pressure increase and voiding during phases III-IV.

**Figure 3:**
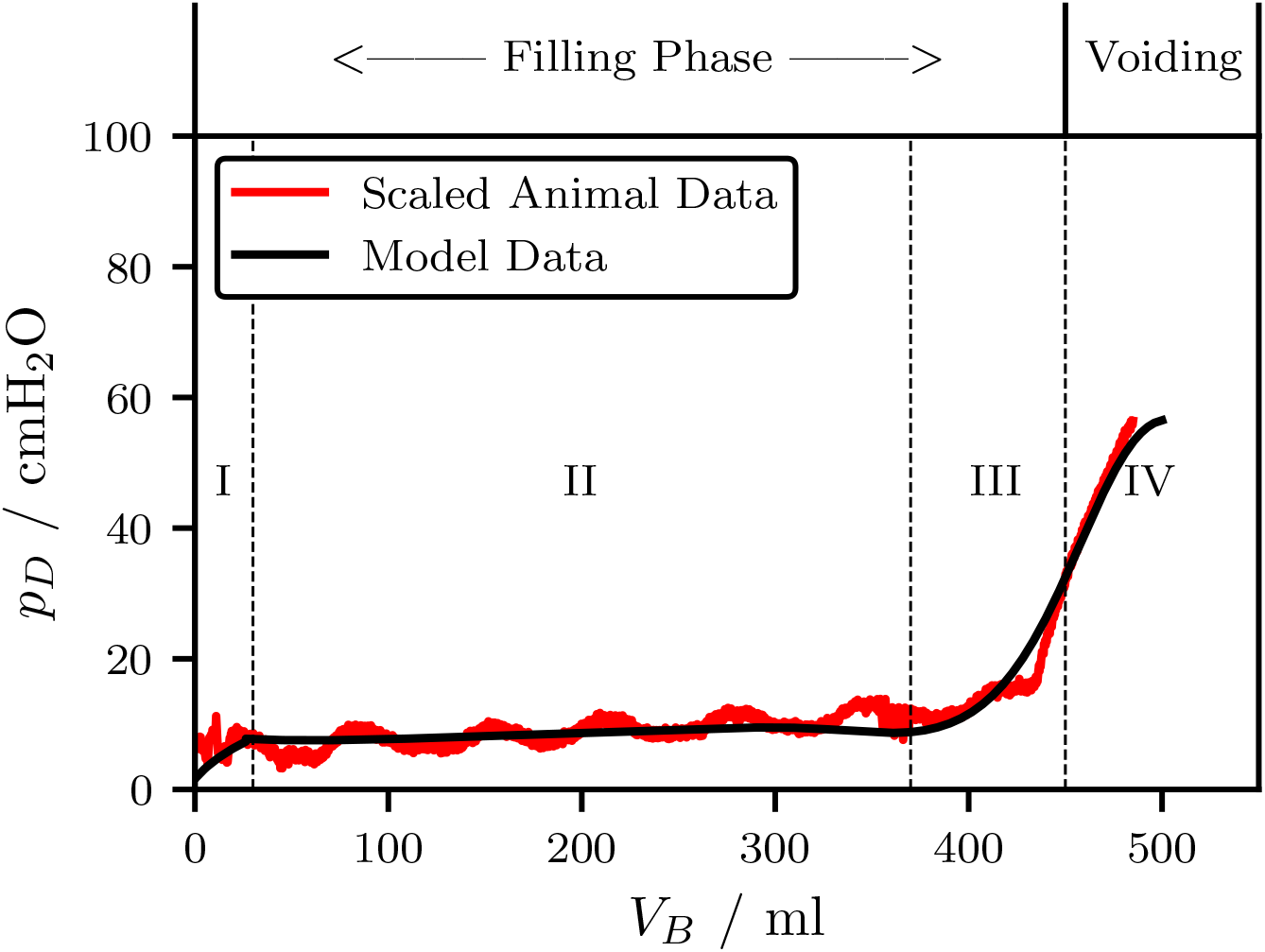
Pressure-volume relationship for scaled animal data and LUT model. The relationship aligns with an ideal human cystometrogram with distinct phases: an initial pressure rise (Phase I), bladder compliance (Phase II), and gradual pressure increase with voiding (Phases III-IV). The scaled animal data and model outputs are in strong agreement, as evidenced by a Pearson correlation coefficient of r = 0.93 (p ! 0.001).

The scaled animal data in Figure 3 exhibited high similarity when overlaid on the human model outputs. This suggests the model effectively captures the general trends observed in the animal experiments. Further strengthening this validation, the Pearson correlation coefficient between the non-scaled animal data and the model outputs yielded a statistically significant value of *r* = 0.93 (*p* < 0.001). This high correlation value highlights a strong positive relationship between the model’s predictions and the in-vivo data, providing further confidence in the model’s accuracy. However, the animal data does demonstrate fluctuations during the compliance phase of storage. This could be due to muscle spasms or noise produced in the recording of the small pressure values.

During phase II of Figure 3, the bladder exhibits compliance as it stretches with minimal increase in detrusor pressure. For a bladder to be considered normal, its compliance should be over 40 mL/cmH_2_O [21]. Given the non-deterministic nature of each voiding cycle, a simulation was performed to assess compliance in every filling cycle. The findings revealed that compliance surpassed 40 mL/cmH_2_O across all cycles, indicating a healthy bladder.

### 3.3. Voiding Dynamics

The stochastic nature of the neural model results in a variable interval between voids, requiring the model to run over 50 days to extract the interval from each void cycle. For the purposes of grouping the intervals, it was assumed that night falls between 10pm and 6am, however this will vary depending on the sleep cycle of the individual. Figure 4 displays the distribution of time intervals between voiding events. While the daytime voiding pattern aligns with the expected 5-7 hour interval, the literature suggests that individuals with lower bladder capacities may experience more frequent voiding, approximately every 2-3 hours [22]. This increased frequency is likely due to the reduced bladder capacity, which necessitates more frequent voiding events to maintain continence. Therefore, the observed voiding pattern aligns with a lower bladder capacity. On the other hand, the night voids seem to be shorter than expected, with the model experiencing voiding on some night. Although this is not uncommon, it could indicate that the filling model needs to be revised beyond a singular sine curve.

**Figure 4:**
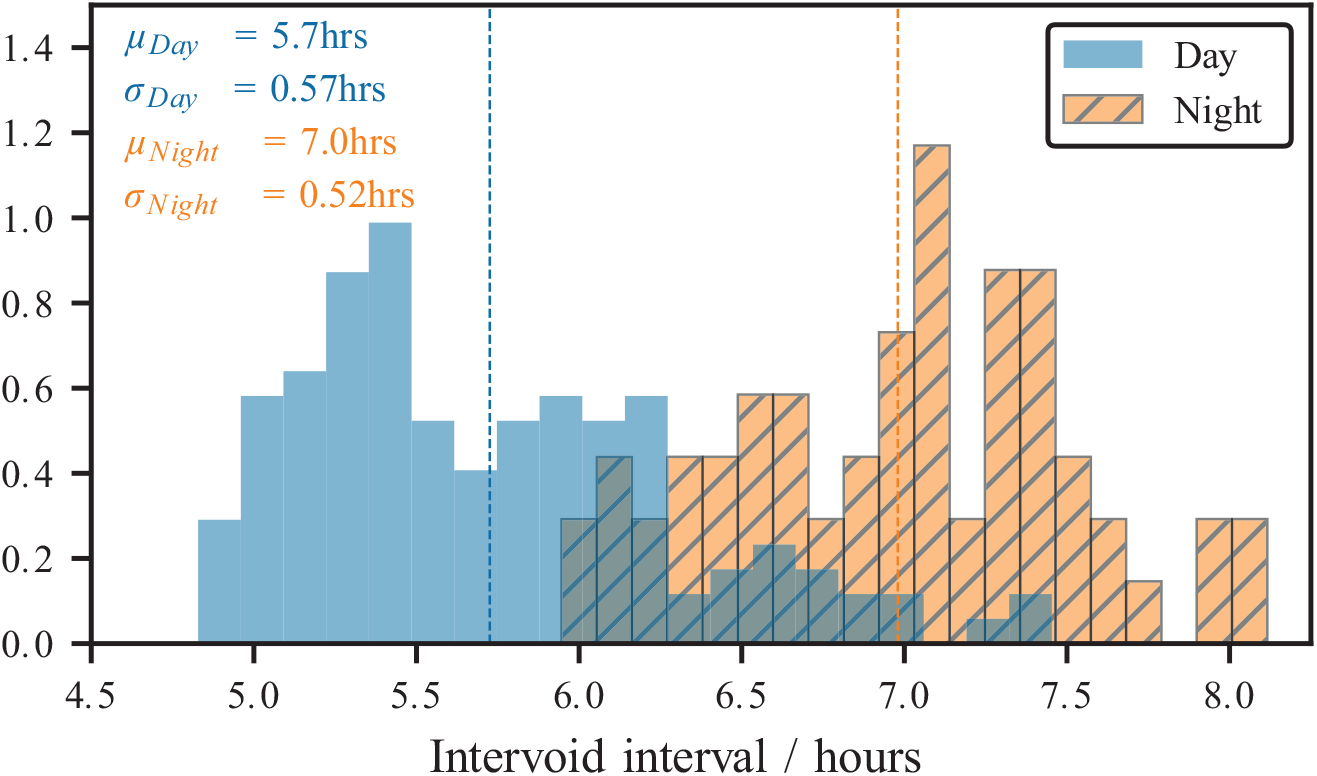
Distribution of Intervoid Intervals. The histogram displays the distribution of times between voiding events. During the day, the model demonstrates a lower mean intervoid interval, indicating more frequent voiding. Conversely, during the night, the model shows less frequent voiding. This behaviour is attributed to the stochastic kidney model integrated into the LUT framework, effectively simulating the natural variations in kidney function over a 24-hour period.

Alongside the assessment of the intervoid interval, the investigation also explored the distribution of voiding durations over the course of the 50-day simulation. The voiding events exhibited a duration of 27.3 ± 0.05 seconds, demonstrating that voiding durations are independent of the preceding bladder filling time. The literature suggests an average voiding duration of 21 ± 13 seconds [23]. While the value in this simulation exceeded the expected duration slightly, it remained within the range established by the literature. The observed difference suggests that modifications to the neural control model, as explored in Subsection 3.1, may be required for improved synchronisation with the voiding dynamics depicted in the current model. The LUT model itself, however, appears to be functioning within a physiologically relevant range.

### 3.4 Noise Sensitivity Analysis

To evaluate the model’s sensitivity to noise, simulations were conducted with varying noise parameter values. The resulting filling rates were analysed and visualised in a box plot (Figure 5). The box plot demonstrates a clear relationship between the noise parameter and the variability of filling rates. As the noise parameter increases, the interquartile range of the filling rates expands, indicating a wider distribution of values. This suggests that the model’s output becomes more sensitive to random fluctuations as the noise level rises. The physiologically relevant range for filling rates is typically defined by a minimum inflow of around 0.3 ml/min, below which oliguria may be indicated [24], and a maximum inflow of around 10 ml/min [25], with an average inflow of approximately 1 ml/min [25]. When noise parameters are less than or equal to 1, the average filling rates fall within this physiological range, with minimal outliers. This shows that the model maintains robustness under these conditions. However, when the noise parameter exceeds 1, the distribution of filling rates becomes more skewed, with a higher frequency of extreme values. When the randomly generated noise component is sufficiently negative, the net inflow drops below zero. To maintain biophysical accuracy, the model imposes a minimum inflow of *Q*_in_ = 0. This prevents unrealistic negative inflows from the kidneys, which are not physiologically possible. Additionally, as the frequency of zero inflows increases, the average inflow rate decreases, leading to longer intervoid intervals. This can result in less frequent voiding, which may not accurately reflect physiological behaviour. While lower noise reduces this effect, it comes at the cost of increased model determinism. So, even though the model occasionally has periods of non-filling at higher noise values (0.5 ≤ *n* ≤ 1), these instances are not a major concern because they occur infrequently enough that the average remains within a safe level. Hence, to ensure model robustness and maintain biophysical accuracy, it is recommended to keep the noise parameter within the range of 0 to 1. If significantly more noise is required, the noise generation method itself should be reconsidered rather than solely adjusting the parameter.

**Figure 5:**
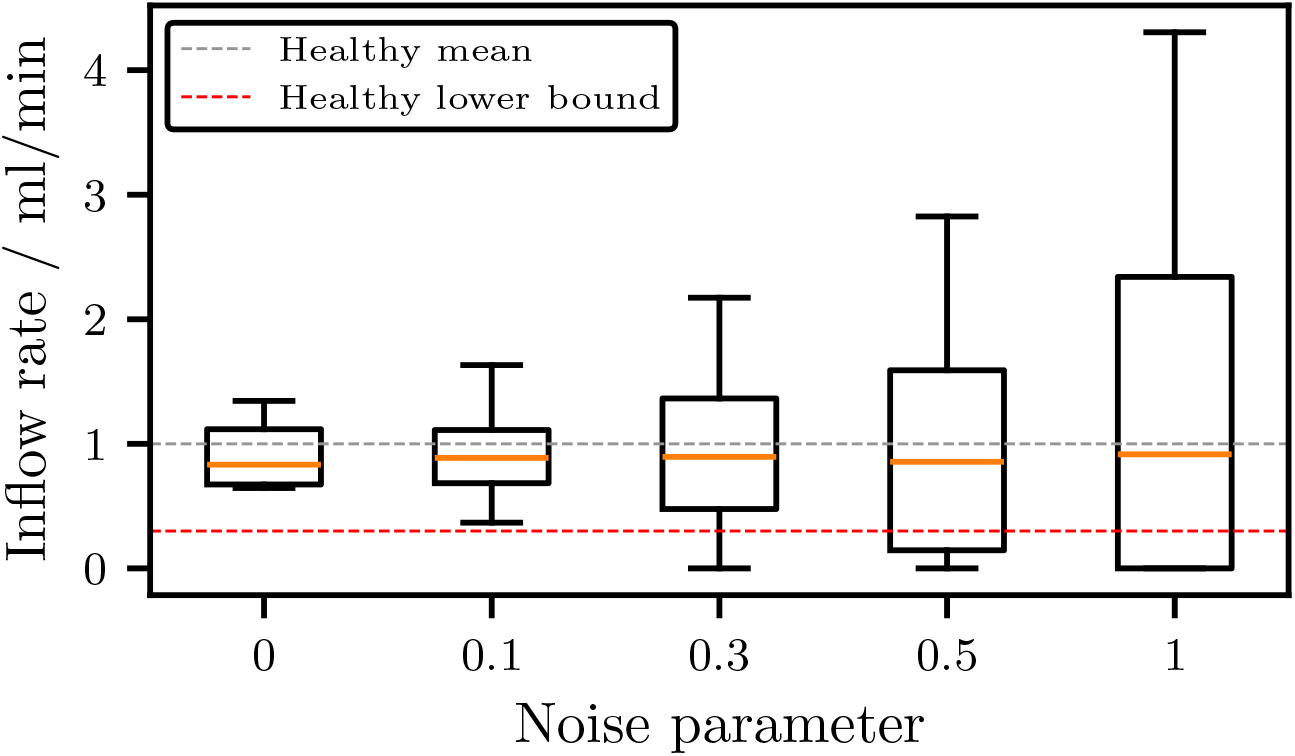
Box plot illustrating the distribution of filling rates under different noise parameter values, with reference lines showing the lower bound of 0.3 ml/min (dashed red) for a healthy filling rate (below which oliguria may be present) and the mean healthy inflow of 1 ml/min (grey).

## 4 Conclusion

This paper presents a novel approach for developing an open-source LUT model incorporating sinusoidal kidney function and noise injection. The model offers the following key attributes: the availability to integrate external neural inputs, smooth neural model representation, variable kidney function, and code accessibility. The model accurately represents the dynamic filling rate and unpredictable behaviour of the biological system, as validated against established physiological ranges and metrics. The model demonstrated robustness by maintaining physiologically healthy filling rates across varying levels of noise within the recommended range. This highlights the resilience of the model to random fluctuations and its ability to provide accurate predictions under different conditions. By introducing random fluctuations, the model can better realistically simulate the natural variations in physiological processes. This feature can lead to a more accurate representation of bladder function and improve the predictive capabilities of the model.

This work contributes to the field of in-silico medicine by providing a valuable tool for studying and understanding bladder function. The availability of an open-source bladder model serves research, education, and clinical applications. In its present form, the model is well suited to analyse pathological conditions associated with physiological or structural changes to the bladder. These include prevalent, life-altering conditions such as Urinary Tract Infection [26], Interstitial Cystitis [27] and obstructive Urinary Retention [28] [29], all of which would benefit considerably from an open-source computational framework for further analysis. From the point of view of future work, there is considerable potential benefit to the integration of this model with a simulation of its neural control system. The research team has recently adapted a modified version of this system which successfully captures the intricate neural interactions underpinning the micturition cycle [30] which they used to analyse the system-level effects of neuromodulation. Were the model outlined here to be integrated with this neural-network, the resulting work could be used to analyse a number of open problems within the lower-urinary tract-pathology literature.

These include the as-of-yet unknown mechanism behind the therapeutic effects of Transcutaneous Tibial Nerve Stimulation [31], or an in-depth exploration of the system level changes that occur as a result of overactive [32], or underactive bladder [33]. The therapeutic value of answering these questions should not be understated. As such, future work will aim to merge these two models to produce a detailed framework that will allow for both bladder-specific, and system-level simulation of pathologies and potential solutions.

## Author Contributions

EL - Conceptualization, Methodology, Software, Validation, Visualization, Writing; AM-T - Project Administration, Supervision, Writing - Review & Editing; MJ - Data Curation. AE - Data Curation; KN - Project Administration, Supervision, Writing - Review & Editing.

All authors gave final approval for publication and agreed to be held accountable for the work performed therein.

